# Stress-induced formation of cell wall-deficient cells in filamentous actinomycetes

**DOI:** 10.1101/094037

**Authors:** K. Ramijan, E. Ultee, J. Willemse, A.J. Wondergem, D. Heinrich, A. Briegel, G.P. van Wezel, D. Claessen

## Abstract

The cell wall is a shape-defining structure that envelopes almost all bacteria. One of its main functions is to serve as a protection barrier to environmental stresses. Bacteria can be forced in a cell wall-deficient state under highly specialized conditions, which are invariably aimed at interrupting cell wall synthesis. Therefore, the relevance of such cells has remained obscure. Here we show that many filamentous actinomycetes have a natural ability to generate a new, cell wall-deficient cell type in response to hyperosmotic stress, which we call S-cells. This wall-deficient state is transient, as S-cells are able to switch to the canonical mycelial mode-of-growth. Remarkably, prolonged exposure of S-cells to hyperosmotic stress yielded variants that are able to proliferate indefinitely without their cell wall. This is the first report that demonstrates the formation of wall-deficient cells as a natural adaptation strategy and their potential transition into stable wall-less forms solely caused by prolonged exposure to osmotic stress. Given that actinomycetes are potent antibiotic producers, our work also provides important insights into how biosynthetic gene clusters and resistance determinants may disseminate into the environment.

## INTRODUCTION

All free-living bacteria are challenged by constant changes in their environment, and their survival depends on the ability to adapt to sudden exposure to stressful conditions. For instance, soil bacteria can encounter rapid osmotic fluctuations caused by rain, flooding, or desiccation. Bacterial cells typically respond to osmotic changes by rapidly modulating the osmotic potential within the cell, either by importing or exporting ions and compatible solutes^1^. While these responses typically occur immediately after cells have been exposed to the changed environment, they are also able to tune the expression of metabolic pathways or critical enzymes^2^.

How such osmotic changes affect cellular morphology is not well known. The cells’ shape is largely dictated by the cell wall, which is a highly dynamic structure that acts as the main barrier that provides osmotic protection^3^. The synthesis of its major constituent, peptidoglycan (PG), involves the activity of large protein complexes that cooperatively build and incorporate new PG precursors into the growing glycan strands at the cell surface^4–7^ These strands are then cross-linked to form a single, giant sacculus that envelops the cell^8^. The sites for the incorporation of new PG is a major difference between the planktonic firmicutes that grow by extension of the lateral wall, and Actinobacteria, which grow via apical extension and thereby incorporating new PG at the cell poles^9, 10^.

Actinobacteria display a wide diversity of morphologies, including cocci (*Rhodococcus*), rods (*Mycobacterium* and *Corynebacterium*) and mycelia (*Streptomyces* and *Kitasatospora*), or even multiple shapes (*Arthrobacter*)^11, 12^. Species belonging to these genera are able to change their morphology to adapt to extreme environments. For example, *Rhodococcus* species that are commonly found in arid environments are able to adapt to desiccation by modulating their lipid content and form short-fragmented cells^13^. *Arthrobacter* species also exhibit high resistance to desiccation and cold stresses. Upon hyperosmotic stress, these cells can modulate the synthesis of osmoprotectants and switch between rod-shaped and myceloid cells^12^.

While the cell wall is considered an essential component of virtually all bacteria, most species can be manipulated under laboratory conditions to produce so-called L-forms that are able to propagate without their wall^14–17^ Typically, L-forms are generated by exposing walled bacteria to high levels of lysozyme combined with antibiotics that target cell wall synthesis in media containing high levels of osmolytes^18, 19^ Stable L-forms that can propagate indefinitely without the cell wall require two mutations that fall in separate classes ^18^. The first class of mutations leads to an increase in membrane synthesis, either directly by increasing fatty acid biosynthesis, or indirectly by reducing cell wall synthesis^20^. The second class of mutations reduces oxidative damage caused by reactive oxygen species, which are detrimental to proliferation of L-forms^21^. Notably, proliferation of L-forms is independent of the FtsZ-based division machinery^15, 22^ Instead, their proliferation can be explained solely by biophysical processes, in which an imbalance between the cell surface area to volume ratio leads to spontaneous blebbing and the subsequent generation of progeny cells^20^. Such a purely biophysical mechanism of L-form proliferation is not species-specific. This observation has led to the hypothesis that early life forms propagated in a similar fashion well before the cell wall had evolved^15, 20, 23^. Whether L-forms have functional relevance in modern bacteria, however, is unclear.

Here we present evidence that many filamentous actinobacteria have a natural ability to extrude cell wall-deficient (CWD) cells when exposed to high levels of osmolytes. These newly-identified cells, which we call S-cells, synthesize PG precursors and are able to switch to the canonical mycelial mode-of-growth. Remarkably, upon prolonged exposure to hyperosmotic stress conditions, S-cells can acquire mutations that enable them to proliferate in the CWD state as so-called S-forms, which are morphologically similar to L-forms but not originating from walled cells exposed to cell wall-targeting agents. These results demonstrate that the extrusion of S-cells and their transition into proliferating S-forms is a natural adaptation strategy in filamentous actinobacteria, solely caused by prolonged exposure to osmotic stress.

## RESULTS

### Hyperosmotic stress drives the formation of cell wall-deficient cells

Recent work suggests that hyperosmotic stress conditions affects apical growth in streptomycetes^24^ Consistent with these observations, we noticed that growth was progressively disturbed in the filamentous actinomycete *Kitasatospora viridifaciens*, when increasing amounts of sucrose were added to the medium (Fig. 1A). In liquid cultures containing more than 0.5 M sucrose, initiation of growth was delayed by at least 5 h compared to media with low levels of sucrose. A similar retardation in growth was observed on solid medium supplemented with high levels of osmolytes, evident from the size decrease of colonies (Fig. 1B, C). On average, their size decreased from 12.8 mm^2^ to 1.4 mm^2^ after 7 days of growth. Notably, the high osmolarity also reduced the number of colony forming units (CFU) by 33%, from 9.3×10^8^ CFU ml^−1^ to 6.1×10^8^ CFU ml^−1^. In order to study the morphological changes accompanying this growth reduction, we stained the mycelium after 48 h of growth with the membrane dye FM5-95 and the DNA stain SYTO-9 (Fig. 1D, E).

**Figure 1.**
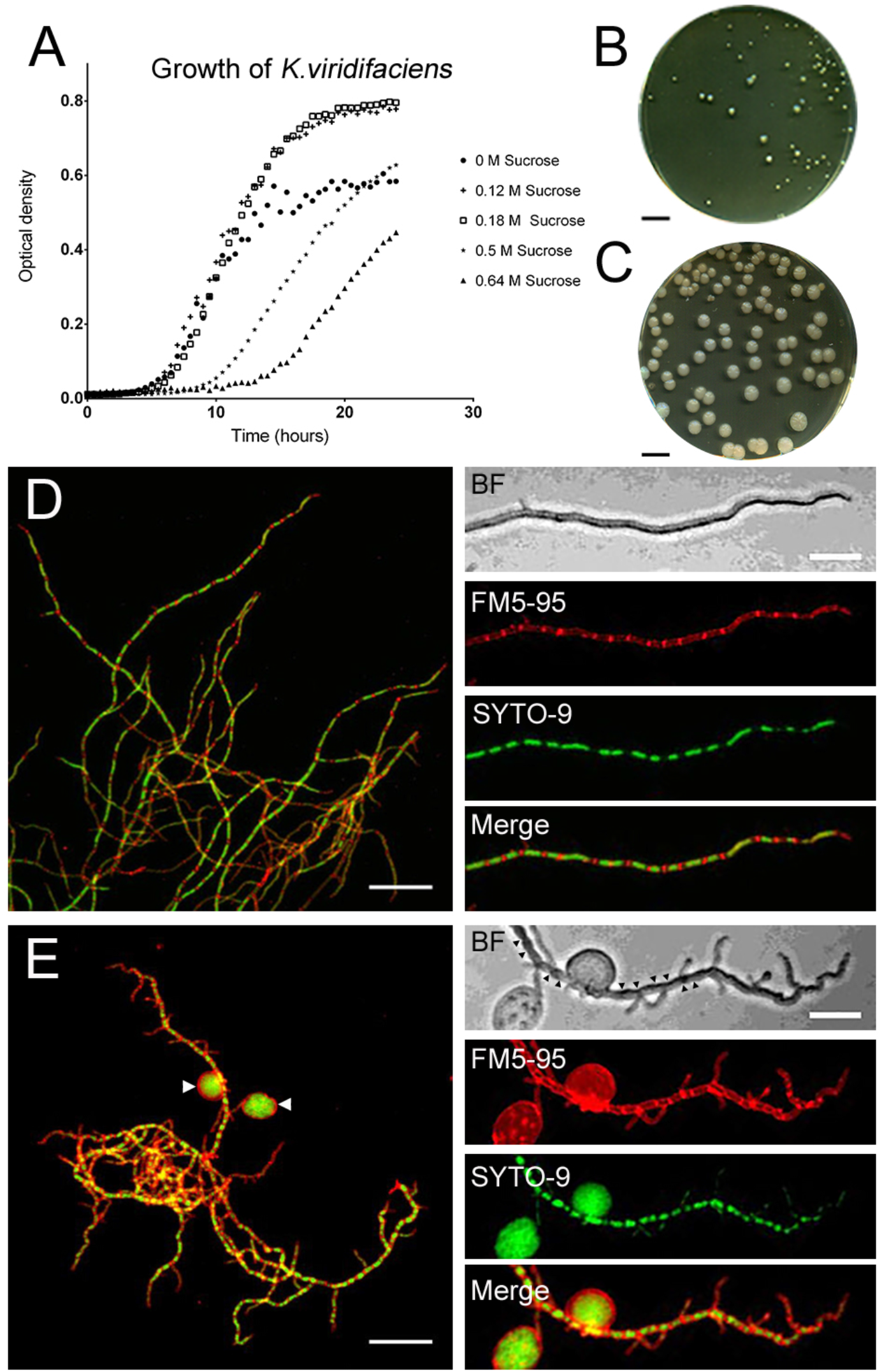
High levels of sucrose affect growth and morphology of *K. viridifaciens*. (A) Growth curves of *K. viridifaciens* in LPB medium supplemented with increasing amounts of sucrose. High levels of osmolytes reduce the number and size of colonies (B) in comparison to media without osmolytes (C). Mycelial morphology *K. viridifaciens* grown in LPB without sucrose (D) and with 0.64M of sucrose (E). Mycelium was stained with FM5-95 and SYTO-9 to visualize membranes and DNA, respectively. Please note the S-cells (arrowheads in E) and indentations along the cylindrical part of the hypha (black arrowheads in BF panel) formed in medium containing high levels of sucrose. Scale bars represent 10 mm (B, C), 10 μm (left panels in D, E) and 20 μm (magnified section in D and E).

The high levels of osmolytes had a dramatic effect on mycelial morphology. The hyphae showed indentations along the cylindrical part of the leading hyphae, reminiscent of initiation of sporulation (see BF panel in Fig. 1E). In addition, the branching frequency increased by more than three-fold in the presence of high levels of osmolytes (Extended Data Table 1 and 2, Student’s T-test, P-value = 0,0010). Additionally, we noticed that these stressed hyphae contained an excess of membrane (compare FM5-95 panels in Fig. 1D, E). The proportion of the hyphae that were stained with FM5-95 increased from 10% to 21% in the presence of 0.64 M Sucrose (Extended Data Table 1 and 2, Student’s T-test, *P*-value < 0,0001). Simultaneously, the average surface area occupied by the nucleoid decreased from 2.59 μm^2^ to 1.83 μm^2^ (Extended Data Table 1 and 2, Student’s T-test, *P*-value = 0.0074). Most strikingly, we observed large DNA-containing vesicles surrounding the mycelial networks (see arrowheads in Fig. 1E). High levels of NaCl had a similar effect on growth and morphology (Fig. S1). *K. viridifaciens* was no longer able to grow when the NaCl concentration was increased to more than 0.6 M (not shown). These results together indicate that upon osmotic stress, the hyphae form a previously uncharacterized cell type, which we hereinafter will refer to as S-cells, for stress-induced cells.

To distinguish S-cells from other cell wall-deficient (CWD) variants of *K. viridifaciens*, we compared them to fresh protoplasts and L-form cells obtained after classical induction with high levels of lysozyme and penicillin G (see Materials and Methods). Size measurements from 2D images revealed that S-cells had an average surface area of 20.73 μm^2^ and were considerably larger than protoplasts and L-forms, which had an average surface area of 4.01 μm^2^ and 7.06 μm^2^, respectively. Vancomycin-BODIPY staining (van^FL^, Fig. S2A) revealed a heterogeneous pattern of nascent PG synthesis in these cells, while in L-forms mostly detached wall material was observed. By contrast, no staining was detected when freshly prepared protoplasts were used (Fig. S2A). When protoplasts were maintained in LPB for 48 hours, their average surface area increased to 7.49+2.21μm^2^, which is considerably smaller than that of S-cells (Extended Data Table 3). Furthermore, protoplasts regenerated a more uniform cell wall while S-cells showed a disordered, non-uniform pattern of cell-wall assembly, whereby wall material was sometimes found to be detached from the cell surface (Fig. S2B, Table Extended Data Table 3).

### Formation of S-cells is common in natural isolates

To see how widespread the formation of S-cells is among natural isolates, we screened our collection of filamentous actinomycetes, obtained from the Himalaya and Qinling mountains^25^, using *Streptomyces coelicolor, Streptomyces lividans, Streptomyces griseus*, and *Streptomyces venezuelae* as the reference strains We used a cut-off diameter of 2 μm to distinguish small S-cells from spores. Spherical cells, similar to S-cells were evident in hyperosmotic media in *S. venezuelae* and in 7 out of the 96 wild isolates (Fig. S3A). The cells were variable in size within the same strains and between strains (Fig. 2A, Extended Data Table 4) and showed differences in the organization of their DNA (Fig. 2A). No S-cells were found in *S. coelicolor, S. griseus*, or *S. lividans* under the tested conditions. Phylogenetic analysis based on 16S rRNA (Fig. S3B), or the taxonomic marker gene *ssgB* used for classifying morphologically complex actinomycetes^26^ revealed that the formation of S-cells is common in at least two genera (Fig. 2B). Moreover, the ability to form S-cells was not restricted to strains that sporulate in liquid-grown cultures. This is based on the observation that MBT86, which belongs to the *S. coelicolor* clade and is classified as a non-liquid sporulating strain, also generates S-cells (Fig. 2C). Altogether, these results show the natural ability to generate S-cells is widespread in filamentous actinomycetes.

**Figure 2.**
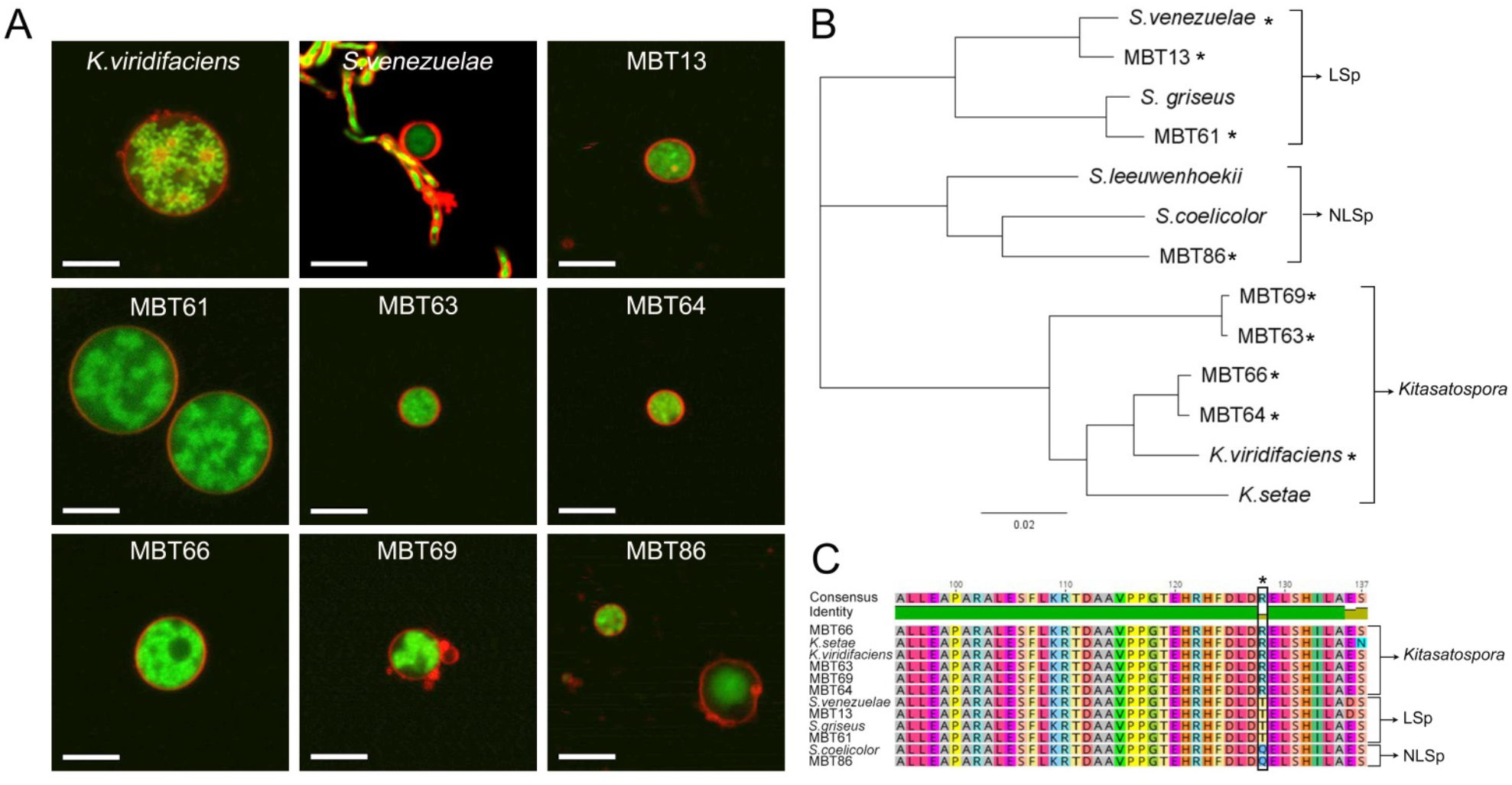
Formation of S-cells is widespread in filamentous actinomycetes. (A) Morphology of S-cells released by *K. viridifaciens, S. venezuelae*, and a number of filamentous actinomycetes from our culture collection (all referred to with the prefix MBT). Cells were stained with FM5-95 (red) and SYTO-9 (green) to visualize membranes and DNA, respectively. (B) Phylogenetic tree of filamentous actinomycetes based on the taxonomic marker *ssgB*. Strains with the ability to form S-cells are indicated with an asterisk (*). *Streptomyces* strains that are able to produce spores in liquid-grown cultures are referred to as LSp (for Liquid Sporulation), while those unable to sporulate in liquid environments are called NLSp (No Liquid Sporulation^26^). This classification is based on amino acid residue 128 in the conserved SsgB protein, which is a threonine (T) or glutamine (Q) for LSp and NLSp strains, respectively. Please note that an arginine (R) is present at this position in all *Kitasatospora* strains (C). Scale bars represent 5 μm

### S-cells are viable cells with the ability to switch to the mycelial mode-of-growth

To determine where S-cells are generated in the hyphae, we performed live imaging of growing germlings of *K. viridifaciens* (Extended Data Video S1). Approximately 7 h after the visible emergence of germ tubes, we detected a transient arrest in tip extension of the leading hypha (Fig. 3A, t=400 mins). Shortly thereafter, small S-cells became visible, which were extruded from the hyphal tip (see arrows in Fig. 3A). These cells rapidly increased in size and number. After 545 min a narrow branch (Fig. 3A arrowhead) was formed in the apical region from which the S-cells were initially extruded. Subapically, other branches became visible approximately 210 minutes after the first appearance of these cells (Extended Data Video S1, t= 770 min). Notably, such branches frequently also extruded S-cells, similarly to the leading hypha (Extended Data Video S2). This showed that S-cells are produced at hyphal tips after apical growth was arrested.

**Figure 3.**
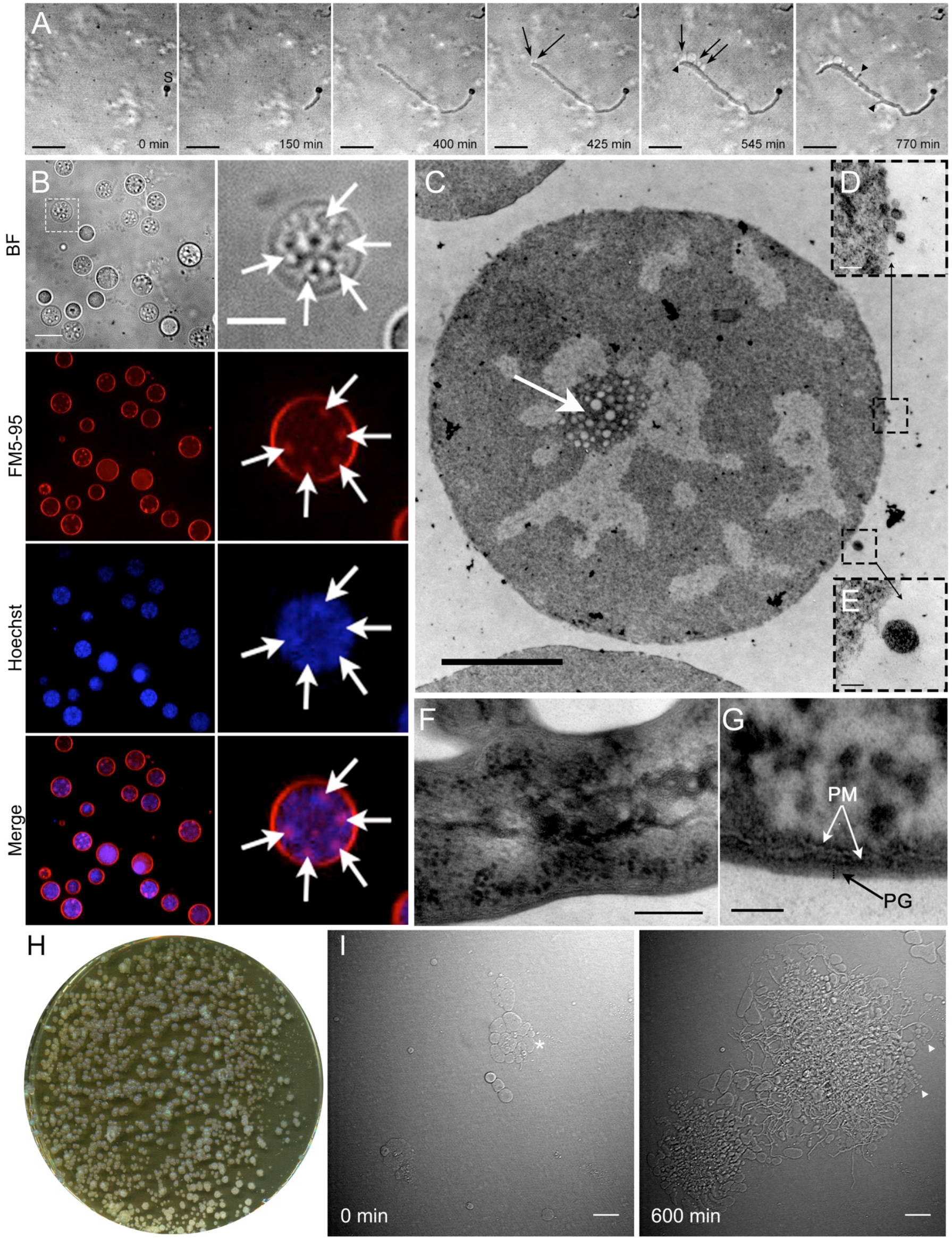
S-cells represent a new cell type with the ability to switch to the mycelial mode-of-growth. (A) Time-lapse microscopy stills showing the extrusion of S-cells (arrows) from hyphal tips. The arrow heads indicate new branches, while “S” designates the germinated spore. Images were taken from Supplementary Video S1 (B) Z-stack projection of filtered S-cells (taken from Supplementary Video 3). Cells were stained with Hoechst and FM5-95 to visualize DNA and membranes, respectively. The intracellular membrane assemblies are indicated with white arrows. (C) Transmission electron micrographs of S-cell reveal the presence of agglomerates of membrane structures (white arrow) in close proximity to the DNA. Contrary to filamentous cells (F, G), S-cells possess a disorganized cell surface (D, E). (H) The S-cells are viable cells with the ability to form mycelial colonies on LPM medium after 7 days of growth. (I) Time-lapse microscopy stills demonstrating the switch of S-cells (asterisk, t=0 min) to filamentous growth. Please note that S-cells are also extruded from newly-formed hyphal tips (arrowheads, t=600 min). Images were taken from Supplementary Video 4. Scale bars represents 10 μm (A, B), 5 μm (magnified section in B), 2 μm (C), 100 nm (D, E), 20 nm (F), 50 nm (G) and 20 μm (I).

Further characterization of S-cells from *K. viridifaciens* revealed that these cells had a granular appearance and membrane assemblies that stained with FM5-95 (Fig. 3B, arrows, Supplementary Video S3). Notably, these assemblies often co-localized with DNA (Fig. 3B, arrows). To study S-cells in more detail, we separated them after 7 days from the mycelia by filtration (see Materials and Methods). In agreement with the previous findings, we also detected agglomerates of membrane assemblies in close proximity of the DNA using electron microscopy analysis (Fig. 3C). Additionally, we noticed that S-cells possessed a disorganized surface, characterized by membrane protrusions that appeared to detach from the S-cells (Fig. 3D, 3E), and an apparent deficiency in normal cell-wall biogenesis (compare to the cell surface of the hypha in Figs. 3F, 3G).

To establish if S-cells were truly viable cells, they were plated onto plates supplemented with sucrose. After 7 days of growth, many mycelial colonies were found; demonstrating that the cells indeed were viable, and that such cell are only transiently CWD (Fig. 3H; ± 1.6×10^4^ CFUs ml^−1^ of the filtered culture). Time-lapse microscopy (Extended Data Video S4) revealed that the cells (Fig. 3I, asterisk) initiated filamentous growth and established mycelial colonies, which, in turn, also extruded new S-cells from the hyphal tips (Fig. 3I, arrowheads). A switch to mycelial growth was also observed when S-cells were inoculated in liquid medium, whether or not the media was supplemented with high levels of sucrose (data not shown). We noticed that the viability of S-cells was reduced by 60% (decreasing from 1.6×10^4^ to 6.7 × 10^3^ CFUs ml^−1^) when these cells were diluted in water before plating. Microscopy analysis indicated that the surviving S-cells were those that showed abundant staining with WGA-Oregon (Fig. S4). Altogether, these results demonstrate that *K. viridifaciens* generates S-cells that synthesize PG and are able to switch to the mycelial mode-of-growth.

### S-cell formation frequently leads to loss of the linear megaplasmid KVP1

When S-cells were allowed to switch to mycelium on MYM medium, we identified many colonies with developmental defects (Fig. 4A). Most obvious was the frequent occurrence of small, brown-colored colonies that failed to produce the white aerial hyphae or the grey-pigmented spores. Non-differentiating colonies are referred to as bald, for the lack of the fluffy aerial hyphae^27^ To test if this aberrant phenotype was maintained in subsequent generations, we selected three of these bald colonies (R3-R5) and two grey-pigmented colonies with a near wild-type morphology (R1 and R2) for further analysis. The progeny of the grey colonies developed similarly to the wild-type strain, and sporulated abundantly after 7 days of growth (Fig. 4B). In contrast, strains R3-R5 failed to sporulate after 7 days of growth. This phenotype is reminiscent of the defective sporulation seen in colonies of *Streptomyces clavuligerus* that have lost the large linear plasmid pSCL4 following protoplast formation and regeneration^28^. Given that *K. viridifaciens* contains a large megaplasmid (KVP1^29^), we reasoned that S-cell formation could increase the frequency of the loss of this plasmid. To test this assumption, we performed quantitative real-time PCR using four genes contained on the megaplasmid (*orf1, parA, tetR*, and *allC*). As a control, we included the two house-keeping genes *infB* and *atpD*, both of which are located on the chromosome, and which encode the translation initiation factor IF-2 and a subunit of the F0F1 ATP synthase, respectively. Detectable amplification of *infB* and *atpD* was seen after 19 PCR cycles in strains R3-R5, which was similar to the wild-type strain (Fig. 4C). The same was true for the KVP1-specific genes *orf1, parA, tetR*, and *allC* in the wild-type strains. However, amplification of these plasmid marker genes was only seen after 30 PCR cycles in strain R3-R5 (Fig. 4D). This demonstrates that the KVP1-specific genes were only present in trace amounts in the R3-R5 strains (at least 10^4^ times less abundant than the chromosomal genes *infB* and *atpD*) (Fig. 4E), which is consistent with loss of KVP1 during formation of S-cells.

**Figure 4.**
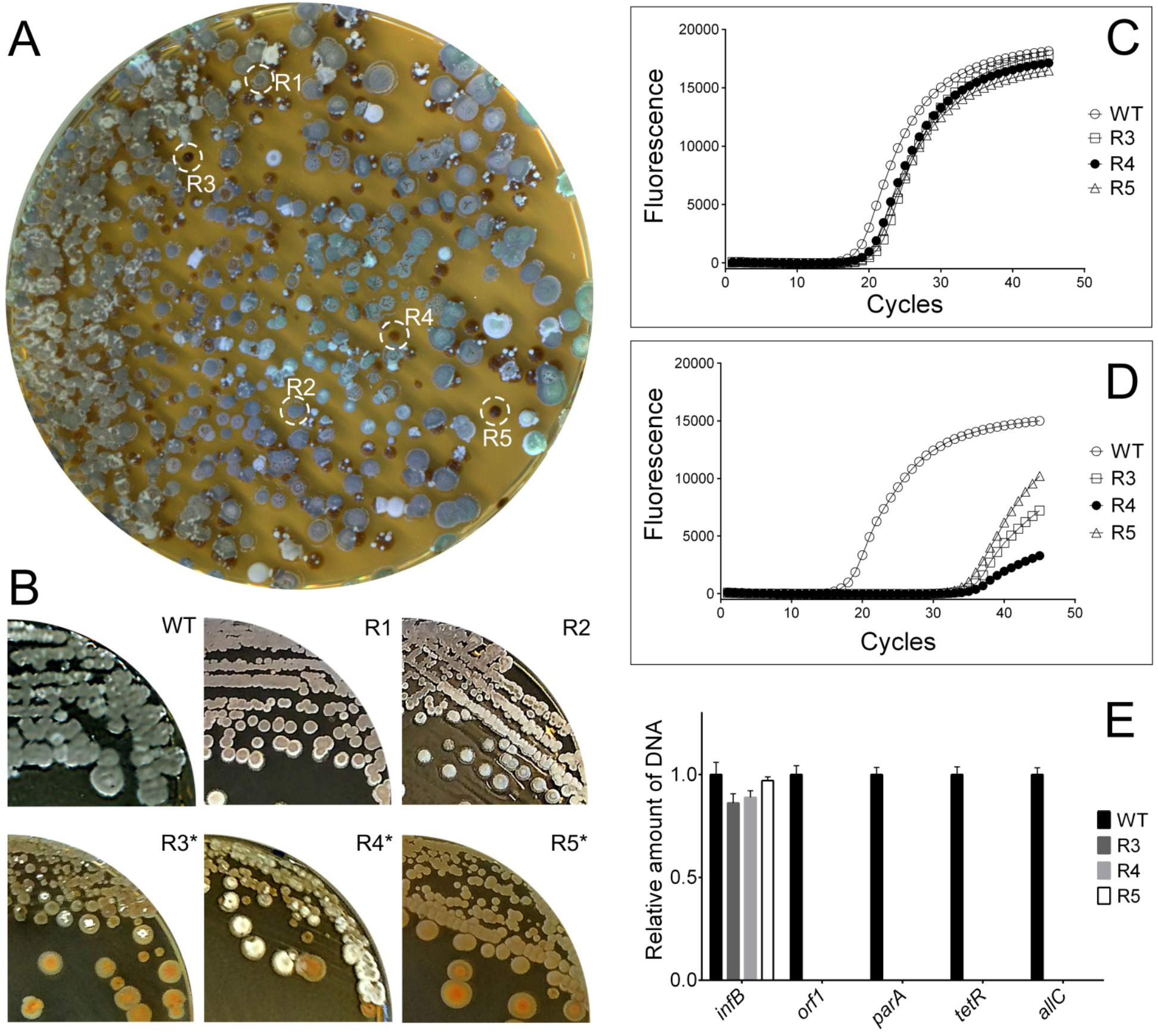
S-cells switching to the mycelial mode-of-growth frequently leads to loss of the megaplasmid KVP1. The switch of S-cells to the mycelial mode-of-growth yields colonies with different morphologies: besides grey-pigmented colonies (R1, R2), colonies are formed that fail to develop efficiently, and which appear whitish or brown (R3-R5). (C) Subculturing of R1 and R2 leads to the formation of grey colonies that appear similar to the wild type, while subculturing of R3, R4 and R5 yield colonies that are unable to form a robust sporulating aerial mycelium (brown and white colonies). Quantitative real time PCR of the *infB* (C) and *allC* (D) genes using gDNA of the wild type and R3-R5 as the template. In all strains, the *infB* gene located on the chromosome is amplified before the 20^th^ cycle. However, the *allC* gene, located on the KVP1 megaplasmid, is amplified in the wild type before the 20^th^ cycle, but in strains R3-R5 after the 30^th^ cycle. (F) Quantitative comparison of the relative abundance of four megaplasmid genes (*orf1, parA, tetR* and *allC*) and the *infB* gene (located on the chromosome) between the wild type and strains R3-R5. The strong reduction in the abundance of the megaplasmid genes are consistent with loss of this plasmid.

### Prolonged hyperosmotic stress is sufficient to convert S-cells into proliferating S-forms

Although the switch to mycelial growth was exclusively observed when young S-cells were cultured in fresh media, we noticed a dramatic change when S-cells had been exposed for prolonged periods to the hyperosmotic stress conditions. In nine out of 15 independent experiments, we found that S-cells switched to mycelial growth, while four times S-cells failed to form a growing culture. Strikingly, however, were the two independent occasions during which S-cells had proliferated in an apparent cell wall-deficient state. On solid medium, these two independent cell lines, called M1 and M2 (for mutants 1 and 2, respectively, see below), formed viscous colonies on LPMA medium, which were similar to those formed by the L-form lineage induced with penicillin and lysozyme, but distinct from the compact colonies formed by the wild-type strain (Fig. 5A). Liquid-grown cultures of M1 and M2 exclusively consisted of CWD cells when sucrose and MgCl_2_ were added (Fig. 5B, BF panels). Staining with WGA-Oregon Green showed these cell lines were indeed cell wall-deficient, as nascent peptidoglycan was mainly found detached from the cell surface (Fig. S5). The spherical cells produced by M1 and M2 were comparable in size to the PenG-induced L-forms. Further microscopic analysis revealed that the cells from M1 and M2 contained inner vesicles (arrowheads in Fig 5B) and tubular protrusions emerging from the cell surface (Fig. 5B, inlay). The vast majority of cells contained DNA, although some empty vesicles were also evident in M1 and M2 (Fig. 5B, asterisks). Time-lapse microscopy revealed that both strains proliferated, whereby smaller progeny cells were released following deformation of the mother cell membrane by either vesiculation (Fig. 5C, taken from Extended Data Video S5), blebbing (Fig. 5D, taken from Extended Data Video S6) or tubulation (Fig. 5E, taken from Extended Data Video S7). Altogether these results inferred that strains M1 and M2 closely resemble the previously described L-forms, both in the inability to regain a cell wall as well as the ability to proliferate in the cell-wall-less state. However, instead of originating from prolonged exposure to antibiotic and/or lysozyme treatment, they originate from osmotically stress-induced cells, and we therefore will call these cells S-forms.

**Figure 5.**
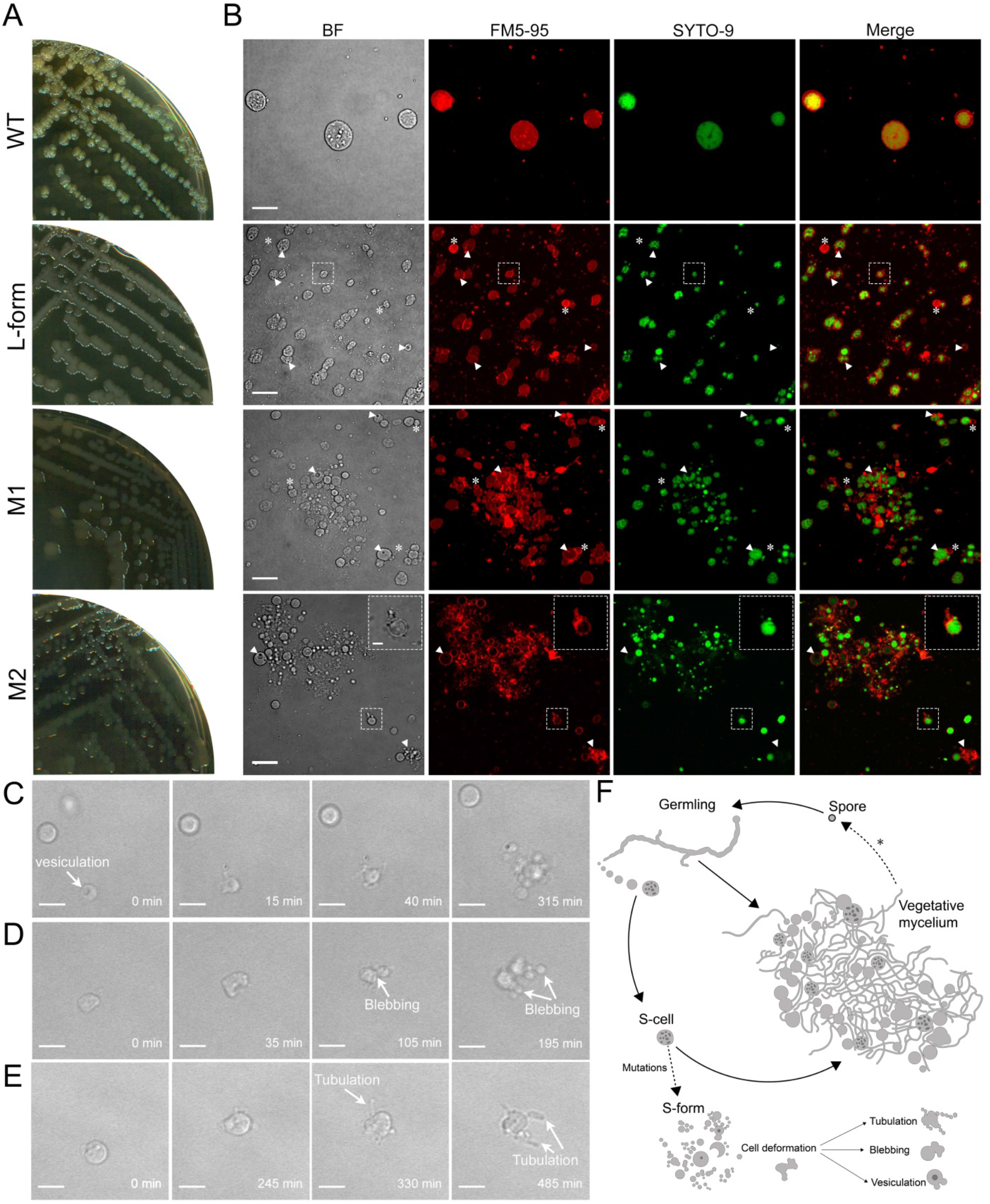
Hyperosmotic stress conditions are sufficient to obtain strains that are able to proliferate without the cell wall. (A) Morphology of colonies of the *K. viridifaciens* wild-type strain, the PenG-induced L-form strain and strains M1 and M2 on LPMA medium. Please note that the wild-type strain forms compact, yellowish mycelial colonies, while the other strains form mucoid green colonies. (B) Morphology of S-cells in comparison to cells of the PenG-induced L-form strain and strains M1 and M2 grown for 48 hours in LPB medium. Cells were stained with FM5-95 and SYTO-9 to visualize membranes and DNA, respectively. Please note that the morphology of cells of strains M1 and M2 is similar to the PenG-induced L-forms. The arrowheads indicate intracellular vesicles, while empty vesicles are indicated with an asterisk. The inlay in M2 shows a proliferation-associated tubulation event. (C-E) Frames from time-lapse microscopy show L-form-like proliferation involving (C) vesiculation, (D) blebbing, and (E) membrane tubulation. (F) Formation of S-cells strains upon prolonged exposure to hyperosmotic stress. Germination of spores under hyperosmotic stress conditions generates germlings, which are able to extrude S-cells. These S-cells are able to switch to the mycelial mode-of-growth, or sporadically acquire mutations that allow them to proliferate like L-forms, which is characterized by tubulation, blebbing or vesiculation. Scale bar represents 10 μm (B), 2 μm (inlay panel B) or 5 μm (C, D, E).

The low frequency at which S-form cells formed made us wonder whether M1 and M2 had acquired mutations that enabled these strains to proliferate without a proper cell wall. Real-time qPCR studies revealed that M1 and M2, but also the PenG-induced L-form cell line, had lost the megaplasmid (Fig. S6). However, loss of the megaplasmid is not sufficient to drive the transition from S-cells to S-forms, as strains R3-R5, all of which had lost the KVP1 megaplasmid, formed mycelia extruding S-cells under hyperosmotic stress conditions (data not shown). Single nucleotide polymorphism (SNP) analysis following whole genome sequencing showed that M1 and M2 had acquired several other mutations (Extended table 5 and 6). Interestingly, both strains carried a mutation in the gene B0Q63_RS21920, which encodes a putative metal ABC transporter. Transporters are often used to cope with osmotic stress conditions^30^. We also identified mutations in the PenG-induced L-form strain (Extended table 7). These mutations, however, differed from those observed in the S-form strains M1 and M2. Notably, the mutations in the PenG-induced L-form appeared to directly relate to cell wall biogenesis, for example in the case of the mutation in *uppP*. The encoded protein is involved in the recycling pathway of the carrier lipid undecaprenyl phosphate (B0Q63_RS22750), which transports glycan biosynthetic intermediates for cell wall synthesis. Altogether, these results demonstrate that prolonged exposures to hyperosmotic stress conditions are apparently sufficient to convert a bacterium into an S-form strain that proliferates without the cell wall.

## DISCUSSION

Filamentous actinomycetes have been intensely studied for more than 50 years as a model for bacterial development. Here, we provide compelling evidence that S-cells represent a natural and previously unnoticed developmental stage in these organisms when they are exposed to hyperosmotic stress conditions (Fig. 5F). These S-cells are extruded from the hyphal tips, they contain DNA and are viable with the ability to grow into mycelial colonies. Furthermore, upon prolonged exposure to hyperosmotic stress, S-cells may also accumulate mutations that enable them to efficiently proliferate in the wall-deficient state we have dubbed S-forms. Our data show that these S-forms can simply emerge as the product of prolonged exposure of cells to hyperosmotic conditions, without directly requiring cell wall-targeting agents. This work provides compelling evidence that such cells have an ecological relevance.

Environmental fluctuations can dramatically influence the availability of water in ecosystems and present osmotic shock conditions to organisms. For instance, microorganisms living in hyperarid regions or hypersaline aquatic environments are frequently exposed to desiccation or hypertonicity^31^. Also, microbes in snow and ice habitats experience low water availability and hypersaline or hyper-acidic environments^32^ Bacteria can adapt to these fluctuations by modulating fatty acid synthesis, accumulating or synthesizing osmoprotectants, protecting their DNA, and secreting extracellular polymeric substance^31, 33^.

Here, we focused on the adaptation of filamentous actinomycetes, which are common in any soil, to extended periods of hyperosmotic stress. As expected, we detected that these bacteria increased the amount of membrane in the hyphae and condensed their nucleoids. Surprisingly though was the extrusion of S-cells. Together with sporulation and the recently discovered explorative mode-of-growth^34^, the ability to form S-cells extends the repertoire by which filamentous actinomycetes can thrive in changing environments. In addition to switching to mycelial growth, these S-cells can have multiple fates. As these cells are wall-deficient, they are prone to lysis due to influx of water. Indeed, exposure to water leads to a steep decline in their ability to outgrow into colonies. However, even when S-cells lyse, the DNA cargo will be released into the environment. Given the large number of biosynthetic gene clusters (BGCs) that are present in the genomes of filamentous actinomycetes, including their resistance determinants, this release of DNA may be a significant, and previously unknown mechanism by which resistance genes are spread. In contrast to releasing DNA into the environment, the S-cells may be able to take up DNA from the environment, similar to other cell wall-deficient cell types such as protoplasts or L-forms^35^. This would enable the cells to acquire genetic information that may help them to overcome the stressful conditions to which they are exposed. In other organisms, this concept has been well characterized. For instance, the bacterium *Bacillus subtilis* becomes naturally competent towards the end of the exponential growth phase^36^. This allows the cells to pick up DNA from the environment, with the prospect of withstanding the harsh conditions and improving the likelihood of survival. Likewise, competence of *Streptococcus pneumoniae* is promoted by exposure to antibiotics that target DNA replication^37^ This, in turn, enables the uptake of foreign DNA (e.g. genes conferring antibiotic resistance). As such, maximizing survival by DNA uptake is a proven strategy.

Our work shows that S-cells are extruded from hyphal tips into the environment, coinciding with an arrest in tip growth. Following their release, the extruding hypha reinitiates growth, indicating that the extrusion process occurs in a manner that apparently is not lethal for the filament from which the cells are released. Tip growth in filamentous actinomycetes is coordinated by the polarisome complex, of which the DivIVA protein is a crucial member^38^. Recent work revealed that hyperosmotic stress has a dramatic effect on the polar growth machinery. Following osmotic upshift experiments, tip growth is arrested, followed by relocation of the apical growth machinery to subapical sites. As a consequence, lateral branches emerge from the leading hyphae^24^. We hypothesize that an imbalance between cell wall synthesis and cell wall turnover could locally lead to changes in the thickness or structure of the cell wall, allowing S-cells to escape from the sacculus.

### Hyperosmotic stress-induced formation of S-forms

L-forms have been studied for many decades, and only recently are we beginning to understand their exciting biology, especially due to ground-breaking work from the Errington lab. L-form cells have been artificially generated from many different bacteria in many laboratories, invariably aimed at targeting the biosynthesis pathway of the cell wall. To that end, cells are typically exposed to high levels of antibiotics, either or not combined with lysozyme treatment^18, 23^. Our work expands on this research by providing for the first-time evidence that CWD strains can emerge solely by exposure to hyperosmotic stress conditions and implies an environmental relevance of this cell type. A crucial and limiting step in the formation of L-forms in *B. subtilis*, as well as in other bacteria, is the escape of a protoplast from the cell-wall sacculus. This process requires lytic activity, which usually comes from lysozyme activity^39^ Our data show that actinomycetes have a natural ability to release such CWD cells when exposed to hyperosmotic conditions. Under prolonged exposure to osmotic stress, some cells are able to acquire mutations allowing these cells to propagate as S-forms.

In line with these findings, recent work shows that *B. subtilis* and *S. aureus* both are able to convert to wall-deficient cells. This has been shown in an animal infection model as well as in macrophages, where lysozyme activity from the host converts walled bacteria into CWD cells^39^. Collectively, these results indicate that cell wall-deficient cells represent an adaptive morphology allowing cells to overcome environmental challenges, such as antibiotic treatment or hyperosmotic stress conditions.

In summary, our work provides evidence for a new, cell wall-deficient cell type in the biology of filamentous actinomycetes. It further expands the large diversity in bacterial cell types, and the plasticity that microorganisms employ to handle environmental stresses. It remains to be elucidated how the ability to form S-cells improves fitness in these filamentous actinomycetes, and how this morphogenetic switch is regulated.

## MATERIALS AND METHODS

### Strains and media

Bacterial strains used in this study are shown in Extended Data Table 8. To obtain sporulating cultures, *Streptomyces* and *Kitasatospora* species were grown at 30°C for 4 days on MYM medium^40^. To support growth of CWD cells, strains were grown on solid medium L-Phase Medium (LPMA), containing 0.5% glucose, 0.5% yeast extract, 0.5% peptone, 20% sucrose, 0.01% MgSO4·7H_2_O, 0.75% Iberian agar (all w/v). After autoclaving, the medium was supplemented with MgCl2 (final concentration of 25 mM) and 5% (v/v) horse serum.

L-Phase Broth (LPB) was used as liquid medium to support growth of wall-deficient cells. LPB contains 0.15% yeast extract, 0.25% bacto-peptone, 0.15% oxoid malt extract, 0. 5% glucose, 0.64 M sucrose, 1.5% oxoid tryptic soy broth powder (all w/v) and 25 mM MgCl2. To test the effect of different sucrose concentrations on mycelial growth and the formation of S-cells, the amount of sucrose in LPB was changed to obtain final concentrations of 0.0, 0.12, 0.18, 0.50 and 0.64 M. The influence of sodium chloride as an osmolyte was analysed by replacing sucrose with NaCl. 50 ml cultures were inoculated with 10^6^ spores ml^−1^ and grown in 250 ml flasks. Cultures were incubated at 30°C, while shaking at 100 rpm.

To prepare protoplasts of *K. viridifaciens*, the wild-type strain was grown for 48 hours in a mixture of TSBS and YEME (1: 1 v/v) supplemented with 5 mM MgCl_2_ and 0.5% glycine. Protoplasts were prepared as described^41^, with the difference that 10 mg ml^−1^ lysozyme solution was used for three hours. Freshly-made protoplast were diluted and immediately used for fluorescence microscopy.

### Optical density measurements

The growth of *K. viridifaciens* was monitored with the Bioscreen C reader system (Oy Growth Curves AB Ltd). To this end, aliquots of 100 μl of LPB medium with different concentrations of sucrose were added to each well of the honeycomb microplate and inoculated with 10^6^ spores ml^−1^. Growth was monitored for 24 hours at 30°C, while shaking continuously at medium speed. The OD wide band was measured every 30 min and corrected for the absorbance of liquid medium without inoculum. In total, five replicate cultures were used for each osmolyte concentration. The effect of sodium chloride as osmolyte was tested using the same procedure, with the differences that the final volume of the cultures was 300 μl, and the experiment was run for 96 hours.

### Quantification of the number and size of colonies

Serial dilutions of *K. viridifaciens* spores were plated in triplicates in LPMA (high osmolarity) and LPMA without sucrose, MgCl_2_ and horse serum (low osmolarity). After 7 days of incubation at 30°C the number of colonies was counted to determine the CFU ml^−1^. Quantification of the surface area of colonies was done with FIJI^42^

### Screening for strains with the ability to release S-cells

To identify strains that are able to release S-cells, strains from an in-house culture collection 25 were initially grown in flat-bottom polysterene 96-well plates, of which each well contained 200 μl LPB medium and 5 μl of spores. The 96-well plate was sealed with parafilm and incubated at 30°C for 7 days. The cultures were then analysed with light microscopy, and strains with the ability to release S-cells with a diameter larger 2 μm were selected. The selected strains were then grown in 250 mL flasks containing 50 mL LPB medium (10^6^ spores ml^−1^) at 30°C while shaking at 100 rpm. After 7 days, aliquots of 50 μl of the bacterial cultures were fluorescently stained with SYTO-9 and FM5-95. The surface area of the S-cells was determined in FIJI^42^ Assuming circularity of these cells, the corresponding diameter D was then calculated as D = 2 * [SQRT(area/π)].

### Filtration of S-cells from *K. viridifaciens*

50 ml LPB cultures of *K. viridifaciens*, inoculated with 10^6^ spores ml^−1^, were grown for 2 or 7 days at 30°C in an orbital shaker at 100 rpm. To separate the S-cells from the mycelium, the cultures were passed through a sterile filter made from an EcoCloth™ wiper. A subsequent filtration step was done by passing the S-cells through a 5 μm Isopore™ membrane filter. The filtered vesicles were centrifuged at 1,000 rpm for 40 mins, after which the supernatant was carefully removed with a 10 mL pipette to avoid disturbance of the S-cells.

### Viability and subculturing of S-cells from *K. viridifaciens*

To verify the viability of S-cells, the filtered cells were incubated in 10 mg ml^−1^ lysozyme solution for 3 hours at 30°C, while shaking at 100 rpm to remove residual hyphal fragments. The filtered S-cells were then centrifuged at 1,000 rpm for 40 mins and resuspended in 1 volume of fresh LPB. Serial dilutions of the S-cells in LPB or water were then plated, in triplicate, on LPMA or MYM medium. The plates were grown for 7 days at 30°C and the CFU values were determined for each treatment.

### Generation of the PenG-induced L-form cell line

Generation of the *K. viridifaciens* L-form lineage was performed by inoculating the wild-type strain in 50 mL LPB medium, supplemented with lysozyme and/or penicillin G, in 250 mL flasks in an orbital shaker at 100 rpm. Every week, 1 mL of this culture was transferred to fresh LPB medium according to the cultivation regime previously described^19^ After the 8^th^ subculture, the inducers were removed from the cultivation medium and the obtained lineage did not revert back to the walled state on LPMA plates or in LPB medium. A single colony obtained after the 8^th^ subculture was designated as PenG-induced L-forms.

### Phylogenetic analysis

The 16S rRNA sequences from strains of the in-house culture collection were previously determined^25^. Homologues of *ssgB* in these strains were identified by BLAST analysis using the *ssgB* sequence from *S. coelicolor* (SCO1541) as the input. For the *Streptomyces* and *Kitasatospora* strains whose genome sequence was not available, the *ssgB* sequence was obtained by PCR with the *ssgB* consensus primers (Extended Data Table 9). Geneious 9.1.7 was used to make alignments of *ssgB* and 16S rRNA, and for constructing neighbour-joining trees.

### Quantitative real time PCR

Filtered S-cells were allowed to regenerate on MYM medium, from which three regenerated bald colonies (R3, R4, and R5) were selected. After two rounds of growth on MYM, bald colonies of the three strains were grown in TSBS for 2 days at 30°C, and genomic DNA was isolated from these strains as described^41^. Primers were designed to amplify the *infB* (BOQ63_RS18295) and *atpD* (BOQ63_RS18295) genes located in the chromosome, and four genes located on the KVP1 megaplasmid: *allC* (BOQ63_RS01235), *tetR* (BOQ63_RS09230), *parA* (BOQ63_RS03875) and *orf1* (BOQ63_RS04285) (Extended Data Table 9). The PCR reactions were performed in triplicate in accordance with the manufacturer’s instructions, using 5 ng of DNA, 5% DMSO and the iTaq Universal SYBR Green Supermix Mix (Bio-Rad). Quantitative real time PCR was performed using a CFX96 Touch Real-Time PCR Detection System (Bio-Rad). To normalize the relative amount of DNA, the wild-type strain was used as a control, using the *atpD* gene as a reference.

### Isolation of the hyperosmotic stress-induced S-form cell lines M1 and M2

Fifteen replicate cultures of *K. viridifaciens* were grown for 7 days in LPB medium. After filtration, the S-cells were transferred to fresh LPB medium. The cultures that had not switched to mycelium after 3 days of cultivation were kept for further analysis. Two cultures turned dark green after 7 days, which after inspection with light microscopy contained proliferating S-form cells. These cell lines were named M1 and M2.

### Microscopy

Bright field images were taken with the Zeiss Axio Lab A1 upright Microscope, equipped with an Axiocam MRc with a resolution of 64.5 nm/pixel.

#### Fluorescence microscopy

Fluorescent dyes (Molecular Probes™) were added directly to 100 μl aliquots of liquid-grown cultures. For visualization of membranes, FM5-95 was used at a final concentration of 0.02 mg ml^−1^. Nucleic acids were stained with 0.5 μM of SYTO-9 or 0.05 mg ml^−1^ of Hoechst 34580. The detection of nascent peptidoglycan was done using 0.02 mg ml^−1^ Wheat Germ Agglutinin (WGA) Oregon Green, or 1 μg ml^−1^ BOPIPY FL vancomycin. Prior to visualization, cells and mycelium were applied on a thin layer of LPMA (without horse serum) covering the glass slides. Confocal microscopy was performed using a Zeiss Axio Imager M1 Microscope. Samples were excited using a 488-nm laser, and fluorescence emissions for SYTO-9, and WGA Oregon Green were monitored in the region between 505–600 nm, while a 560 nm long pass filter was used to detect FM5-95. Detailed fluorescence microscopy pictures (i.e. those in Figures 1D-E, S1C-D, 5B) represent average Z-projections of stacks.

The characterization of the membrane assemblies in S-cells was done on a Nikon Eclipse Ti-E inverted microscope equipped with a confocal spinning disk unit (CSU-X1) operated at 10,000 rpm (Yokogawa, Japan) using a 100x Plan Fluor Lens (Nikon, Japan) and illuminated in bright-field and fluorescence. Samples were excited at wavelengths of 405 nm and 561 nm for Hoechst and FM5-95, respectively. Fluorescence images were created with a 435 nm long pass filter for Hoechst, and 590-650 nm band pass for FM5-95. Z-stacks shown in Extended Data Video S3 were acquired at 0.2 μm intervals using a NI-DAQ controlled Piezo element.

Visualization of stained CWD cells for size measurements were done using the Zeiss Axio Observer Z1 microscope. Aliquots of 100 μl of stained cells were deposited in each well of the ibiTreat μ-slide chamber (ibidi®). Samples were excited with laser light at wavelengths of 488, the green fluorescence (SYTO-9, BODIPY FL vancomycin, WGA-Oregon) images were created with the 505-550 nm band pass, while a 650 nm long pass filter was used to detect FM-595. An average Z-stack projection was used to make Fig. S2A.

#### Time-lapse microscopy

To visualize the emergence of S-cells, spores of *K. viridifaciens* were pre-germinated in TSBS medium for 5 hours. An aliquot of 10 μl of the recovered germlings was placed on the bottom of an ibiTreat 35 mm low imaging dish (ibidi®), after which an LPMA patch was placed on top of the germlings.

To visualize switching of S-cells, these cells were collected after 7 days by filtration from a *K. viridifaciens* LPB liquid-grown culture. A 50 μl aliquot of the filtrate was placed on the bottom of an ibiTreat 35 mm low imaging dish (ibidi®) with a patch of LPMA on top.

To visualize the proliferation of M1 and M2, the strains were grown for 48 hours in LPB. Aliquots of the culture were collected, and centrifuged at 9,000 rpm for 1 min, after which the supernatant was removed, and the cells resuspended in fresh LPB. Serial dilutions of the cells were placed in wells of a ibiTreat μ-slide chamber (ibidi®).

All samples were imaged for ~15 hours using an inverted Zeiss Axio Observer Z1 microscope equipped with a Temp Module S (PECON) stage-top set to 30°C. Z-stacks with a 1 μm spacing were taken every five minutes using a 40x water immersion objective. Average intensity projections of the in-focus frames were used to compile the final movies. Light intensity over time was equalised using the correct bleach plugin of FIJI.

#### Electron microscopy

To visualize the vegetative mycelium of *K. viridifaciens* by transmission electron microscopy (TEM), the strain was grown in TSBS medium for 48 hours. An aliquot of 1.5 ml of the cultures was centrifuged for 10 mins at 1,000 rpm, after which the supernatant was carefully removed with a pipette. The mycelium was washed with 1X PBS prior to fixation with 1.5% glutaraldehyde for one hour at room temperature. The fixed mycelium was centrifuged with 2% low melting point agarose. The solid agarose containing the embedded mycelium was sectioned in 1 mm^3^ blocks, which were post-fixed with 1% osmium tetroxide for one hour. The samples were then dehydrated by passing through an ethanol gradient (70%, 80%, 90% and 100%, 15 min per step). After incubation in 100% ethanol, samples were replaced in propylene oxide for 15 minutes followed by incubation in a mixture of Epon and propylene oxide (1:1) and pure Epon (each step one hour). Finally, the samples were embedded in Epon and sectioned into 70 nm slices, which were placed on 200-mesh copper grids. Samples were stained using uranyl-430 acetate (2%) and lead-citrate (0.4%), if necessary, and imaged at 70 kV in a Jeol 1010 transmission electron microscope.

To image S-cells, a culture of the wild-type *K.viridifaciens* strain that had been grown in LPB medium for 7 days was immediately fixed for one hour with 1.5% glutaraldehyde. Filtered S-cells (see above) were then washed twice with 1X PBS prior to embedding in 2% low melting agarose. A post-fixation step with 1% OsO4 was performed before samples were embedded in Epon and sectioned into 70 nm slices (as described above). Samples were stained using uranyl-430 acetate (2%) and lead-citrate (0.4%), if necessary, and imaged at 70 kV in a Jeol 1010 transmission electron microscope

#### Image analysis

Image analysis was performed using the FIJI software package. To describe the morphological changes during hyperosmotic stress, we compared mycelium grown in LPB with or without 0.64 M of sucrose (i.e. the concentration in LPB medium). After making average Z-stack projections from mycelia, 10 hyphae derived from independent mycelia projections were further analysed. For each hypha, the total length was measured using the segmented line tool and the number of branches emerging from that hypha was counted. The hyphal branching ratio was calculated as the number of branches per μm of leading hypha.

To calculate the surface area occupied by membrane in hyphae either or not exposed to 0.64 M sucrose, we divided the total surface area that stained with FM5-95 by the total surface area of the hypha. FIJI was also used to measure the average surface area of the nucleoid (using SYTO-9 staining) in both growth conditions. Student’s T-tests with two-sample unequal variance were performed to calculate *P*-values and to discriminate between the samples.

To determine the size of cell wall-deficient (CWD) cells, we compared cells of PenG-induced L-form to fresh protoplasts and S-cells, all obtained or prepared after 48 hours of growth. Cells were stained with FM5-95 and SYTO-9 and deposited in the wells of an ibiTreat μ-slide chamber (ibidi®). The size of the spherical was determined as the surface area enclosed by the FM5-95-stained membrane. For the particular case of L-forms, where empty vesicles are frequent, only cells that contained DNA were measured. At least 200 cells of each CWD variant were analysed. Proliferating L-forms in which the mother cell could not be separated from the progeny, were counted as one cell.

### Genome sequencing and SNP analysis

Whole-genome sequencing followed by *de novo* assembly (Illumina and PacBio) and variant calling analyses were performed by BaseClear (Leiden, The Netherlands). The unique mutations were identified by direct comparison to the parental strain *Kitasatospora viridifaciens* DSM40239 (GenBank accession number PRJNA353578^29^). The single and multiple nucleotide variations were identified using a minimum sequencing coverage of 50 and a variant frequency of 70%. To reduce the false positives the initial variation list was filtered, and the genes with unique mutations were further analysed. All variants were verified by sequencing PCR fragments (primer sequence in Extended table S9).

---

**Figure S1. High levels of salt affect growth and morphology of *K. viridifaciens*.** (A) Growth curves of *K. viridifaciens* in LPB medium supplemented with different amounts of NaCl. (B) S-cells (arrowheads) are evident after 96 hours of growth in the presence of 0.3 (middle) or 0.6 M NaCl (bottom). Mycelial morphology of *K. viridifaciens* grown in the presence of 0.6 M NaCl after 48 (C), and 96 hours (D). Mycelium was stained with FM5-95 and SYTO-9 to visualize membranes and DNA, respectively. After 48 hours in the presence of NaCl, only small aggregates of spores and germlings were visible. Pellets obtained after 96 hours showed an excess of membrane and hypercondensation of DNA. Scale bars represent 10 μm (B), or 20 μm (C-D).

**Figure S2. Comparison between different types of cell wall-deficient cells of *K. viridifaciens*.** (A) Morphology of freshly made protoplasts (top panels), PenG-induced L-forms (middle panels) and S-cells (bottom panels). Cells were stained with the membrane dye FM5-95 or fluorescent vancomycin (van^FL^) to detect nascent PG. (B) Morphology of protoplasts (top panels) and S-cells (bottom panels) grown for 48 hours in LPB. Cells were stained with the membrane dye FM5-95 or wheat germ agglutinin (WGA-Oregon) to detect newly synthesized PG. Scale bars represents 10 μm.

**Figure S3. Formation of S-cells is a natural adaptation in filamentous actinomycetes**. (A) Microscopic analysis of strains grown for 7 days in liquid medium containing high levels of osmolytes. Cells were stained with FM5-95 and SYTO-9 to visualize membranes and DNA, respectively. Arrowheads indicate S-cells produced by the different strains. (B) Phylogenetic tree of filamentous actinomycetes based on the 16S rDNA gene. Strains with the ability to form S-cells are indicated with an asterisk (*). Scale bars represents 10 μm.

**Figure S4. The presence of abundant peptidoglycan surrounding S-cells confers resistance to water treatment**. (A) Filtered S-cells were stained with the membrane dye FM5-95 and WGA-Oregon to stain nascent peptidoglycan. The inlay shows an S-cell possessing abundant cell wall material surrounding the cell surface. (B) Morphology of S-cells after exposing them to water. While many cells lyse, some S-cells remain intact, which invariably have abundant cell wall material associated with their cell surface (see inlay). Scale bars represents 10 μm (A-B), and 5 μm (inlays).

**Figure S5. The hyperosmotic stress-induced strains M1 and M2 are cell wall-deficient variants**. Morphology of cells of strains M1 and M2 grown for 48 hours in LPB. Cells were stained with WGA-Oregon to visualize nascent PG. The scale bar represents 10 μm.

**Figure S6. Proliferation in the cell wall-deficient state leads to loss of the megaplasmid KVPI**. Quantitative real time PCR of the *infB* (A) and *allC* (B) genes using gDNA of the wild-type strain, the PenG-induced L-form strain, and strains M1 and M2. In all strains, the *infB* gene located on the chromosome is amplified before the 20^th^ cycle. However, the *allC* gene, located on the KVP1 megaplasmid, is only amplified in the wild type strain, but not in any of the other strains. (F) Quantitative comparison of the relative abundance of four megaplasmid genes (*orf1, parA, tetR* and *allC*) and the *infB* gene (located on the chromosome) between the wild-type strain, the PenG-induced L-form strain, and strains M1 and M2.

**Extended Data Video S1. Apical extrusion of S-cells in *K. viridifaciens*.** S-cells are extruded from the hyphal tip after 425 min, coinciding with a transient arrest in tip growth. After extrusion of S-cells, a new tip is formed in the apical region of the hyphae after 540 min, while subapically new branches become visible after 620 min. The times are indicated in min. The scale bar represents 10 μm.

**Extended Data Video S2. Extrusion of S-cells from branches in *K. viridifaciens*.** S-cells are extruded from the tips of branches that are formed subapically. The times are indicated in min. Scale bar represents 10 μm.

**Extended Data Video S3. S-cells of *K. viridifaciens*** contain DNA and inner membrane assemblies. Z-stack projects of S-cells isolated after 48 hours, which were stained with Hoechst (blue) and FM5-95 (red) to visualize DNA and membranes, respectively. The scale bar represents 10 μm.

**Extended Data Video S4. Switching of S-cells to the mycelial mode-of-growth**. Switching of S-cells on solid LPMA medium yields colonies consisting of both hyphae and S-cells. The times are indicated in min. The scale bar indicates 20 μm.

**Extended Data Video S5. Example of vesiculation during proliferation of strain M2**. Time-lapse microscopy showing proliferation of strain M2 on media containing high levels of sucrose. Please note that vesiculation is evident in some cells. The times are indicated in min. The scale bar indicates 5 μm.

**Extended Data Video S6. Example of blebbing during proliferation of strain M2**. Time-lapse microscopy showing proliferation of strain M2 on media containing high levels of sucrose. Please note that blebbing is evident in some cells. The times are indicated in min. The scale bar indicates 5 μm.

**Extended Data Video S7. Example of membrane tubulation during proliferation of strain M2**. Time-lapse microscopy showing membrane tubulation in strain M2 during proliferation on media containing high levels of sucrose. The times are indicated in min. The scale bar indicates 5 μm.

**Extended Data Table 1.** Image analysis measurements on hyphae formed in the presence of low levels of osmolytes

| Hyphae | Length (µm) | Number of branches | Branching frequency | Membrane fraction | Average nucleoid area (µm<sup>2</sup>) |
| --- | --- | --- | --- | --- | --- |
| 1 | 86,0180 | 1 | 0,01 | 0,10 | 2,53 |
| 2 | 90,377 | 3 | 0,03 | 0,09 | 2,01 |
| 3 | 72,3597 | 7 | 0,10 | 0,10 | 1,90 |
| 4 | 83,9838 | 0 | 0,00 | 0,10 | 3,02 |
| 5 | 79,0436 | 0 | 0,00 | 0,09 | 3,21 |
| 6 | 65,3853 | 1 | 0,02 | 0,17 | 4,18 |
| 7 | 80,7872 | 7 | 0,09 | 0,07 | 2,11 |
| 8 | 86,3086 | 2 | 0,02 | 0,10 | 3,05 |
| 9 | 76,4282 | 3 | 0,04 | 0,08 | 2,14 |
| 10 | 74,1033 | 2 | 0,03 | 0,12 | 1,71 |
| Average | 79,4795 | 2,6 | 0,03 | 0,10 | 2,59 |
| SD | 7,5857 | 2,5 | 0,03 | 0,03 | 0,77 |

**Extended Data Table 2.** Image analysis measurements on hyphae formed in the presence of high levels of osmolytes

| Hyphae | length<br>($\mu\text{m}$) | Number of<br>branches | Branching<br>frequency | Membrane<br>Fraction | Average nucleoid<br>area ($\mu\text{m}^2$) |
| --- | --- | --- | --- | --- | --- |
| 1 | 82,5308 | 11 | 0,13 | 0,22 | 1,35 |
| 2 | 90,9580 | 6 | 0,07 | 0,18 | 2,10 |
| 3 | 88,3428 | 8 | 0,09 | 0,16 | 1,82 |
| 4 | 60,4451 | 12 | 0,20 | 0,29 | 1,86 |
| 5 | 86,0180 | 9 | 0,10 | 0,24 | 2,08 |
| 6 | 76,7188 | 6 | 0,08 | 0,21 | 2,05 |
| 7 | 88,6334 | 6 | 0,07 | 0,22 | 2,34 |
| 8 | 53,4707 | 8 | 0,15 | 0,19 | 1,47 |
| 9 | 69,1631 | 7 | 0,10 | 0,20 | 2,04 |
| 10 | 70,3255 | 2 | 0,03 | 0,21 | 1,15 |
| Average | 76,6606 | 7,5 | 0,10 | 0,21 | 1,83 |
| STDEVA | 12,9242 | 2,8 | 0,05 | 0,03 | 0,38 |

**Extended Data Table 3.** Comparison between *K.viridifaciens* cell wall-deficient cells

| Characteristic | Protoplast | L-form | S-cell |
| --- | --- | --- | --- |
| Origin | Osmoprotective conditions combined with lysozyme treatment | Osmoprotective conditions combined with prolonged exposure to lysozyme and PenG | Osmoprotective conditions |
| Area ($\mu\text{m}^2$) | 4.01±1.93 | 7,06±5.87 | 20.73±11.53 |
| Cell wall | Homogeneous regeneration. Wall material mostly associated with the cell surface | Not uniform, disordered assembly. Wall material often detached from the cell surface | Not uniform, disordered assembly. Wall material sometimes detached from the cell surface |
| Genotype | Wild type | Mutant | Wild-type |

**Extended Data Table 4.**
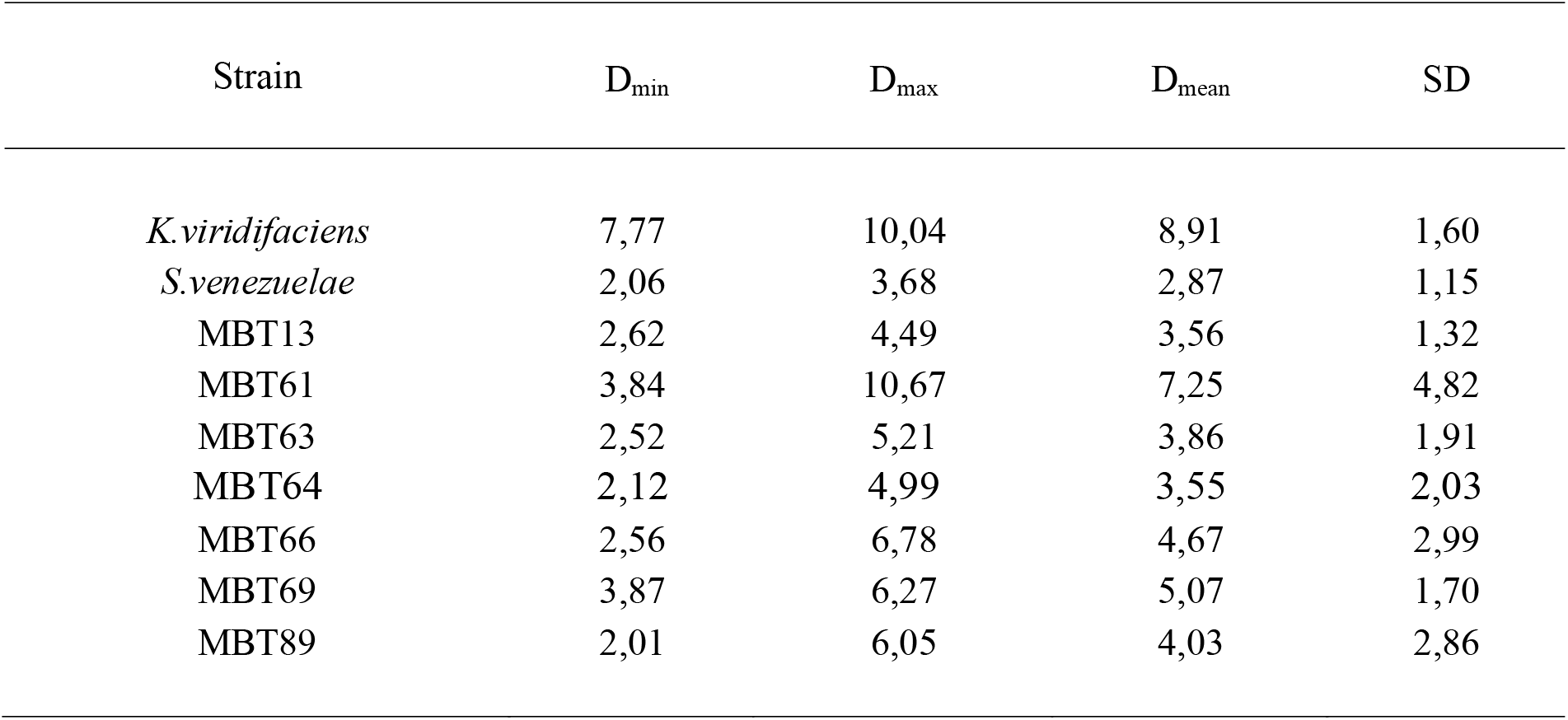
Calculated diameters (D) of S-cells released by different filamentous actinomycetes upon hyperosmotic stress. The diameters are indicated in μm.

| Strain | D<sub>min</sub> | D<sub>max</sub> | D<sub>mean</sub> | SD |
| --- | --- | --- | --- | --- |
| *K.viridifaciens* | 7,77 | 10,04 | 8,91 | 1,60 |
| *S.venezuelae* | 2,06 | 3,68 | 2,87 | 1,15 |
| MBT13 | 2,62 | 4,49 | 3,56 | 1,32 |
| MBT61 | 3,84 | 10,67 | 7,25 | 4,82 |
| MBT63 | 2,52 | 5,21 | 3,86 | 1,91 |
| MBT64 | 2,12 | 4,99 | 3,55 | 2,03 |
| MBT66 | 2,56 | 6,78 | 4,67 | 2,99 |
| MBT69 | 3,87 | 6,27 | 5,07 | 1,70 |
| MBT89 | 2,01 | 6,05 | 4,03 | 2,86 |

**Extended Data Table 5.**
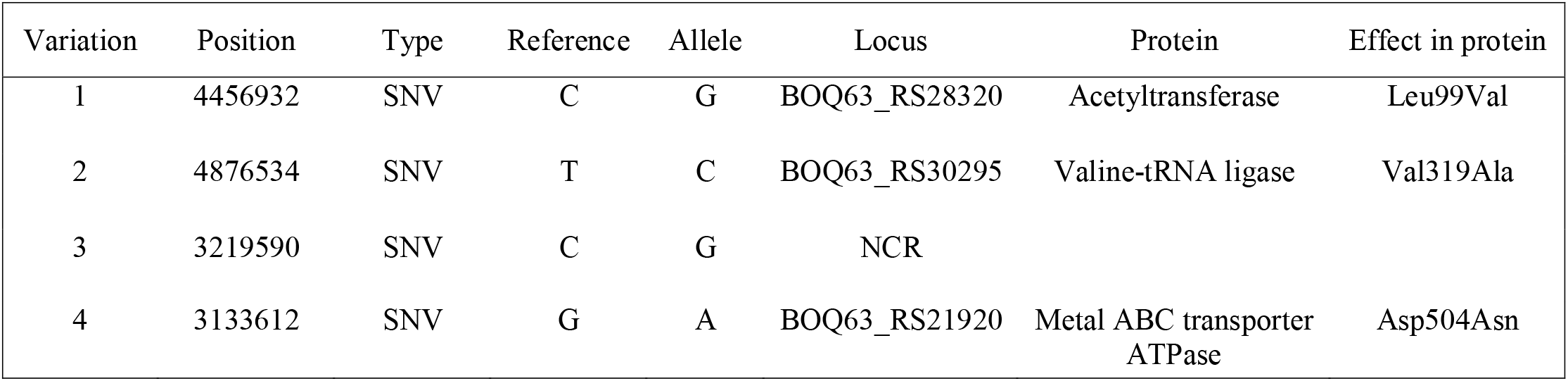
Mutations in the hyperosmotic stress-induced S-form strain M1 SNV: Single Nucleotide Variation

**Extended Data Table 6.** Mutations in the hyperosmotic stress-induced S-form strain M2 SNV: Single Nucleotide Variation, del: deletion, fs: frame shift

| Variation | Position | Type | Reference | Allele | Locus | Protein | Effect in protein |
| --- | --- | --- | --- | --- | --- | --- | --- |
| 1 | 2164717 | SNV | A | G | NCR |  |  |
| 2 | 5054842 | SNV | T | G | BOQ63_RS31145 | XRE family<br>transcriptional<br>regulator | Glu332Ala |
| 3 | 6460621-6460623 | Deletion | CCA | - | BOQ63_RS37840 | Histidine kinase | Thr606del |
| 4 | 3 133753^<br>3133754 | Insertion | - | C | BOQ63_RS21920 | Metal ABC transporter<br>ATPase | Arg553fs |

SNV: Single Nucleotide Variation, del: deletion, fs: frame shift

**Extended Data Table 7.**
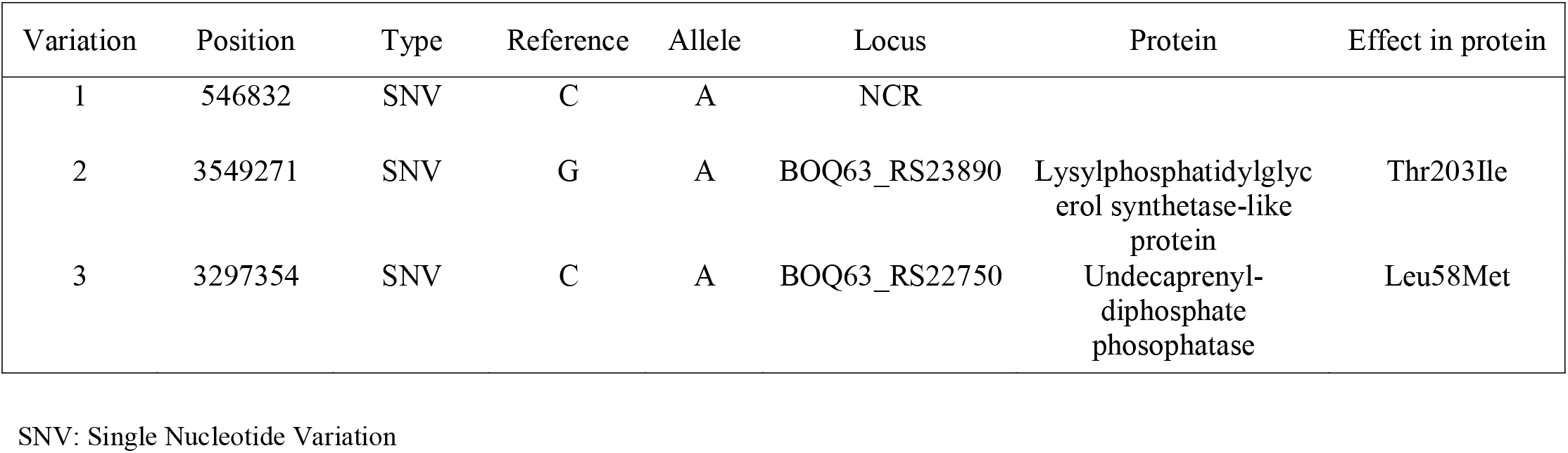
Mutations in the PenG induced L-form SNV: Single Nucleotide Variation

| Variation | Position | Type | Reference | Allele | Locus | Protein | Effect in protein |
| --- | --- | --- | --- | --- | --- | --- | --- |
| 1 | 546832 | SNV | C | A | NCR |  |  |
| 2 | 3549271 | SNV | G | A | BOQ63_RS23890 | Lysylphosphatidylglycerol synthetase-like protein | Thr203Ile |
| 3 | 3297354 | SNV | C | A | BOQ63_RS22750 | Undecaprenyl-diphosphate phosphatase | Leu58Met |

SNV: Single Nucleotide Variation

**Extended Data Table 8.** Strains used in this study

| Strains | Genotype | Reference |
| --- | --- | --- |
| ***Streptomyces/Kitasatospora* strains** |  |  |
| *Streptomyces coelicolor* A3(2) M145 | Wild-type | Lab collection |
| *Streptomyces lividans* 1326 | Wild-type | Lab collection |
| *Streptomyces griseus* | Wild-type | Lab collection |
| *Streptomyces venezuelae* DIVERSA | Wild-type | Lab collection |
| *Kitasatospora viridifaciens* DSM40239 | Wild-type | DSMZ, (Ramijan et al. 2017) |
| *Streptomyces* sp. MBT13 | Wild-type | Lab collection (Zhu et al. 2014) |
| *Streptomyces* sp. MBT61 | Wild-type | Lab collection (Zhu et al. 2014) |
| *Kitasatospora* sp. MBT63 | Wild-type | Lab collection (Girard, 2014) |
| *Kitasatospora* sp. MBT64 | Wild-type | Lab collection (Zhu et al. 2014) |
| *Kitasatospora* sp. MBT66 | Wild-type | Lab collection (Girard, 2014) |
| *Kitasatospora* sp. MBT69 | Wild-type | Lab collection (Zhu et al. 2014) |
| *Streptomyces* sp. MBT86 | Wild-type | Lab collection (Zhu et al. 2014) |
| ***K.viridifaciens* cell wall-deficient strains** |  |  |
| PenG-induced L-form | Mutant | This work |
| Hyperosmotic stress-induced S-form M1 | Mutant | This work |
| Hyperosmotic stress-induced S-form M2 | Mutant | This work |

**Extended Data Table 9.**
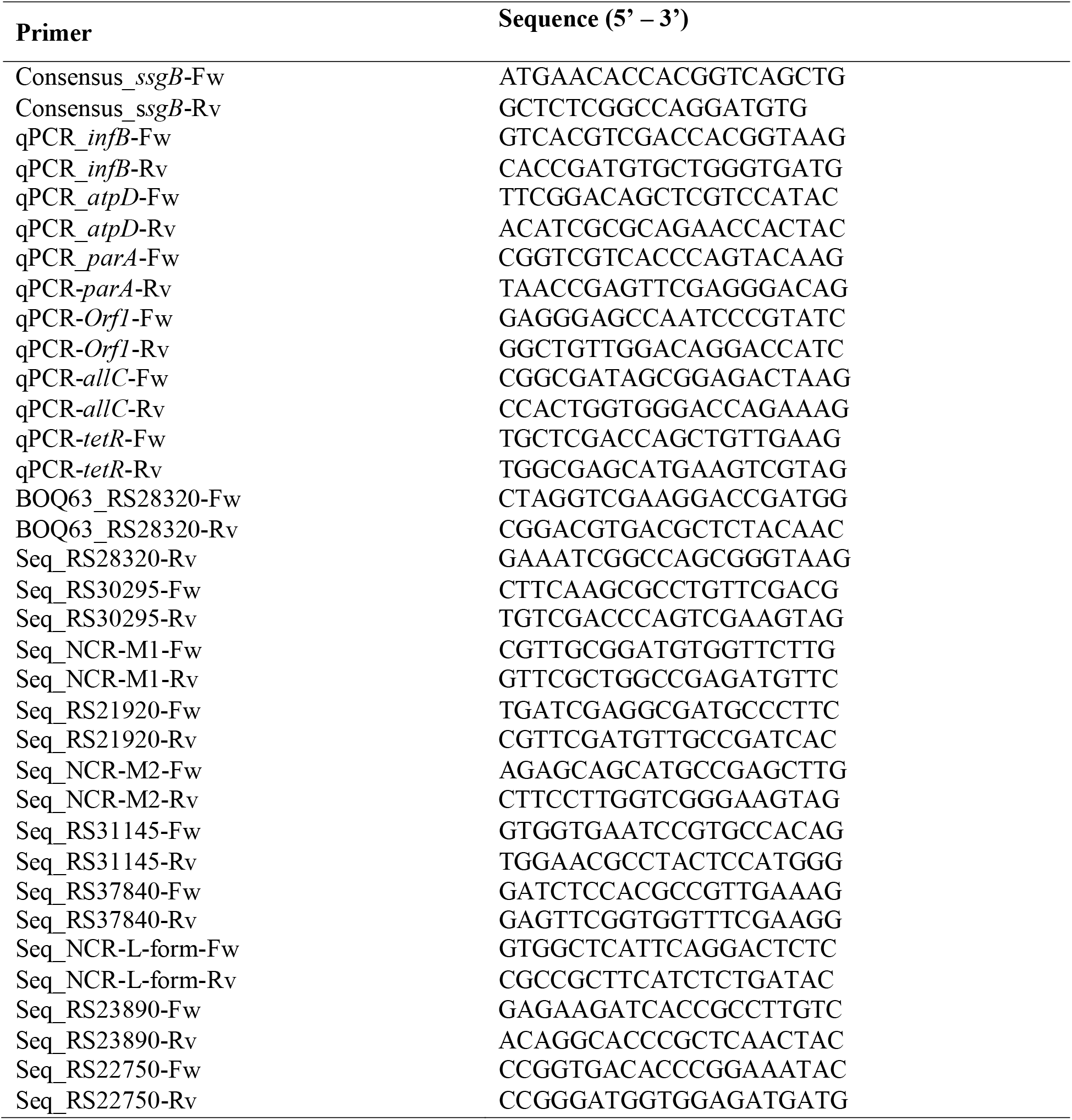
Primers used in this study

